# Climate change vulnerability for species – assessing the assessments

**DOI:** 10.1101/062646

**Authors:** Christopher J. Wheatley, Colin M. Beale, Richard B. Bradbury, James W. Pearce-Higgins, Rob Critchlow, Chris D. Thomas

## Abstract

Climate change vulnerability assessments are commonly used to identify species at risk from global climate change, but the wide range of methodologies available makes it difficult for end users, such as conservation practitioners or policy makers, to decide which method to use as a basis for decision-making. Here, we compare the outputs of 12 such climate change vulnerability assessment methodologies, using both real and simulated species, and we test the methods using historic data for British birds and butterflies (i.e., using historical data to assign risks, and more recent data for validation). Our results highlight considerable inconsistencies in species risk assignment across all the approaches considered and suggest the majority of the frameworks are poor predictors of risk under climate change – two methods performed worse than random. Methods that incorporated historic trend data were the only ones to have any validity at predicting distributional trends in subsequent time periods.

---

Standardised methods of risk assessment are important tools for prioritising adaptive strategies to counter the impacts of climate change, including conservation action for species most likely to face extinction. The IUCN Red List^1,2^ is globally accepted as the method for assessing the vulnerability of species to extinction. However, it has recently been suggested that this process does not adequately identify potential future risk, such as that posed by climate change, as it focuses more on the symptoms of declines than on the underlying causes^3^. Given that global extinction risks^4–6^ are increasing as a consequence of climate change^7, 8^, this could potentially lead to an under-estimate of the risk to species^7^. These concerns have led to the parallel development of a number of risk assessment frameworks^9^, each of which aims to quantify the vulnerability or extinction risk of a species due to climate change.

Each framework draws on different input variables and combines them in different ways, so they are not necessarily interchangeable. To allow for meaningful interpretation of the assessments by conservation practitioners and policy makers, it is necessary to evaluate whether the results of different frameworks are in agreement with one another; and this is currently unknown. If the results of species risk assessments do differ, the choice of framework would affect the perceived vulnerability of different species, hence changing conservation priorities and management actions. It is also unknown whether any of the different assessment frameworks provide a projection of risk that is accurate or realistic. It is important, therefore, that the frameworks should be validated using empirical data on observed changes to the status of species to determine which methods are most appropriate to use.

Climate change vulnerability assessments follow two broad approaches^9^: trait-based and trend-based. Trait-based vulnerability assessment frameworks^10–14^ focus primarily on species traits believed to increase or decrease risk under climate change. These include traditional traits, such as life-history information, but they may also incorporate trait data derived from distributional data (e.g. to estimate thermal limits). In contrast, trend-based correlative frameworks^15–17^ focus primarily on abundance and distribution changes (observed and projected), supplemented by some trait information to inform assessors of the likelihood that projected trends will be realised. Some studies have attempted to combine the two types into hybrid frameworks^18–21^, weighting one set of inputs most heavily or including trend-based data as an optional set of inputs. The ease of applying each of these frameworks depends on the availability of trait, trend and modelled input data for the taxon and region under consideration. In this regard, some frameworks have been developed with specific taxa in mind^10–12,16,19–21^, particularly birds and other vertebrates, while others are generic; and they have been applied to a range of geographic scales (Table 1). However, they can all be scaled up or applied to different taxonomic groups with little or no adjustment.

**Table 1.** Summary vulnerability framework information. Overall vulnerability equation used by each framework, broad methodology type, taxonomic group(s) used to test the framework, and geographic scale at which the framework was tested. The Pearce-Higgins et al. 2015 framework is a simplified version of the Thomas et al. 2011 framework, excluding exacerbating factors and including only trend data.

| General vulnerability equation | Framework | Methodology type | Taxon | Locality |
| --- | --- | --- | --- | --- |
| Exposure x sensitivity | Gardali et al. 2012<sup>12</sup> | Trait | Birds | California State |
|  | Young et al. 2012<sup>18</sup> | Hybrid | Molluscs, Fish, Amphibians, Birds, Mammals | Nevada State |
|  | Moyle et al. 2013<sup>19</sup> | Hybrid | Freshwater fish | California State |
|  | Garnett et al. 2013<sup>21</sup> | Hybrid | Birds | Australia |
|  | Thomas et al. 2011<sup>15</sup> | Trend | Birds, Plants, Invertebrates, | Great Britain |
|  | Pearce-Higgins et al. 2015<sup>17</sup> | Trend | Birds, Plants, Invertebrates | Great Britain |
| Exposure x sensitivity x conservation status | Triviño et al. 2013<sup>16</sup> | Trend | Birds | Iberian Peninsula |
| Exposure x sensitivity x adaptive capacity | Chin et al. 2010<sup>10</sup> | Trait | Chondrichthyan fish | Great Barrier Reef |
|  | Foden et al. 2013<sup>13</sup> | Trait | Birds, Amphibians and Corals | Global |
| Exposure + sensitivity | Barrows et al. 2014<sup>14</sup> | Trait | Plants, Mammals, Reptiles, Birds | Joshua Tree National Park |
|  | Heikkinen et al. 2010<sup>20</sup> | Hybrid | Butterflies | Europe |
| Exposure + sensitivity + adaptive capacity | Arribas et al. 2012<sup>11</sup> | Trait | Water beetles | Iberian Peninsula |

In general, the frameworks attempt to quantify three major components (or some combination thereof) of risk: sensitivity, exposure and adaptive capacity^22, 23^. All approaches, whether trait-or trend-based, explicitly incorporate measures that are intended to represent both species exposure and species sensitivity to climate change (Table 1) but, beyond this, there is little agreement across the frameworks on exactly which measures (input variables) to use. This may arise, in part, because there is limited evidence to identify which traits are most important in determining the sensitivity of a species to climate change^24^ or exactly how climate exposure should be quantified. A range of different inputs are therefore used to assess vulnerability, using a combination of projections from distribution models, population dynamics and life history traits. These amount to 117 specific input variables across the 12 frameworks considered here, of which three-quarters are unique to a single framework; and only 5 of the 117 variables are represented in more than two frameworks (Supplementary Table 1). Ideally, these differences would not matter and each framework would identify the same species as vulnerable, but this should be tested, not assumed. In addition to the variation in input variables used by different frameworks, there is inconsistency in whether inputs are considered measures of sensitivity, exposure or adaptive capacity. For example, metrics of dispersal are treated as sensitivity^12,14,15,18,20^, exposure^19^ or adaptive capacity^10,11,13^ depending on the framework used.

Here, we assess the utility of 12 published frameworks, using some of the best biodiversity data available. Initially, we consider whether the 12 frameworks generate consistent results; i.e. whether the frameworks ‘agree’ on which species are at risk from climate change. We also consider the current Red List assessment approach, without incorporating any future projected declines using bioclimate envelope modelling, and compare the outputs against those from each of the 12 frameworks. We then validate the performance of the 12 different frameworks. We carry out an assessment using each framework based on historic species data and compare the outcomes to subsequent, observed changes in distribution and population. For frameworks that perform well in validation, species that are classified as at risk using historical data are expected to be most likely to have declined since then.

## Results

### Consistency between the results of different vulnerability frameworks

We first assessed risk to 18 data-rich species (11 bird and 7 butterfly) in Great Britain (hereafter ‘exemplar species’, Table 2) using each of the 12 frameworks and a medium emissions scenario. Individual frameworks differed in their risk categories, so we standardised the output from each to a low/medium/high scale (Supplementary Table 2). The results of the assessments were highly variable, with no single exemplar species assigned to the same risk category by all frameworks (Table 2). The majority of species were classified as high risk by at least one assessment (14/18 species); yet only one species (Black Grouse) was classified as high risk by at least half of the frameworks (Table 2). Pairwise Spearman’s rank correlations between frameworks showed poor overall agreement in risk assignment (r_s_ mean = 0.17 ± 0.03, r_s_ median = 0.21).

**Table 2.**
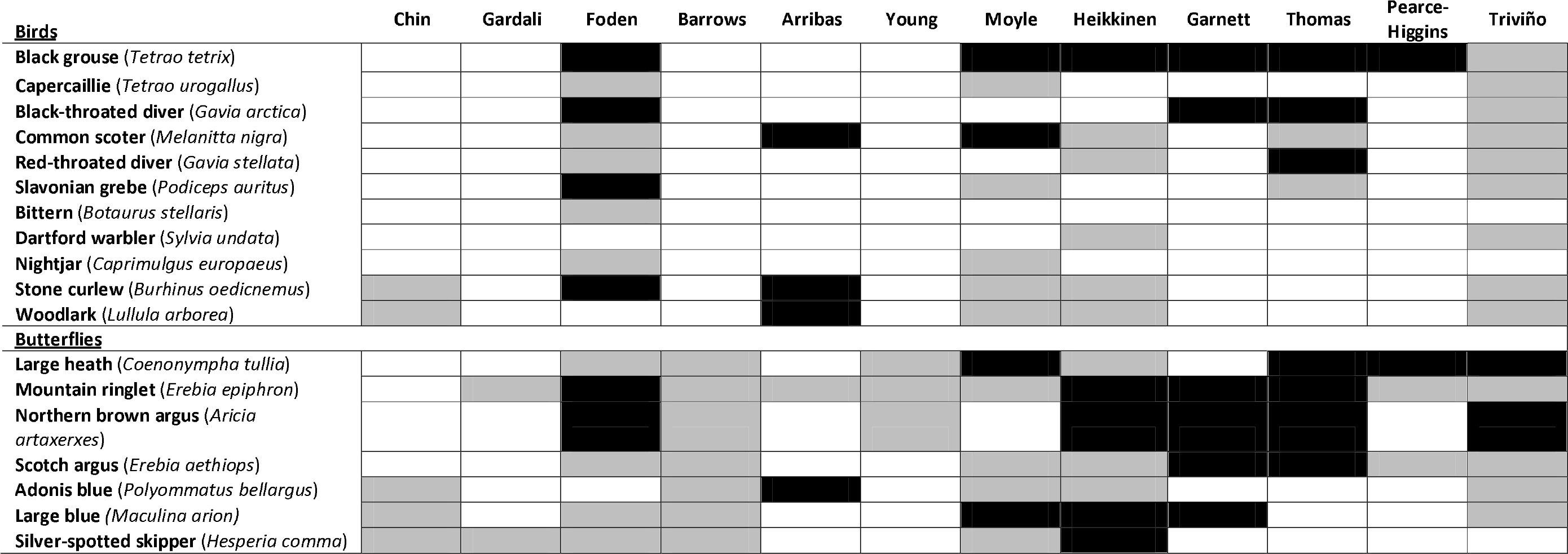
Risk assessment output for exemplar real species. Low (white), Medium (grey) and High (black) risk category outputs for each of the 18 exemplar species assessed using all 12 climate change vulnerability assessment frameworks. Assessments were carried out at the Great Britain scale, based upon contemporary data, with modelled future distributions based upon a medium emission scenario (A1B projection for 2070-2099).

As conservation prioritisation will ultimately concentrate on high risk species, we then focussed only on classification of species in the highest risk category. Inter-rater reliability analysis (for high risk versus low or medium risk) produced a similar pattern to the rank correlation results, with ‘weak’^25^ agreement across frameworks (mean k_PABAK_ = 0.51 ± 0.03, median k_PABAK_ = 0.55). A similar pattern was observed for the exemplar taxa when using a low emissions climate scenario, with only a small number of species changing risk categories between scenarios (Supplementary Table 3). The frameworks also showed poor overall agreement with the Red List assessment (r_s_ mean = −0.28 ± 0.03, r_s_ median = −0.25), and this agreement was not improved when we considered trait-based and trend-based frameworks separately (trait-based: r_s_ mean = −0.39 ± 0.02, trend-based: r_s_ mean = 0.01 ± 0.01).

We further tested the frameworks with an additional 171 British bird and 47 British butterfly species (Supplementary Table 4) for which data were available to model GB distribution changes under a medium emissions climate change scenario. Of these 218 species, 119 were classified as high risk by at least one framework (54%) (Figure 1B), with only 13 species (3 bird and 10 butterfly species) classified into the same risk category by every framework (Supplementary Table 4). Pairwise rank correlations showed poor overall agreement (r_s_ mean = 0.18 ± 0.03, r_s_ median = 0.17), confirming that even with a larger sample of real species with strong correlations between traits, there was little consistency across the frameworks. In addition, inter-rater reliability analysis indicated weak^25^ agreement across frameworks when classifying species as high risk (mean κ_PABAK_ = 0.43 ± 0.03, median K_PABAK_ = 0.61).

**Figure 1.**
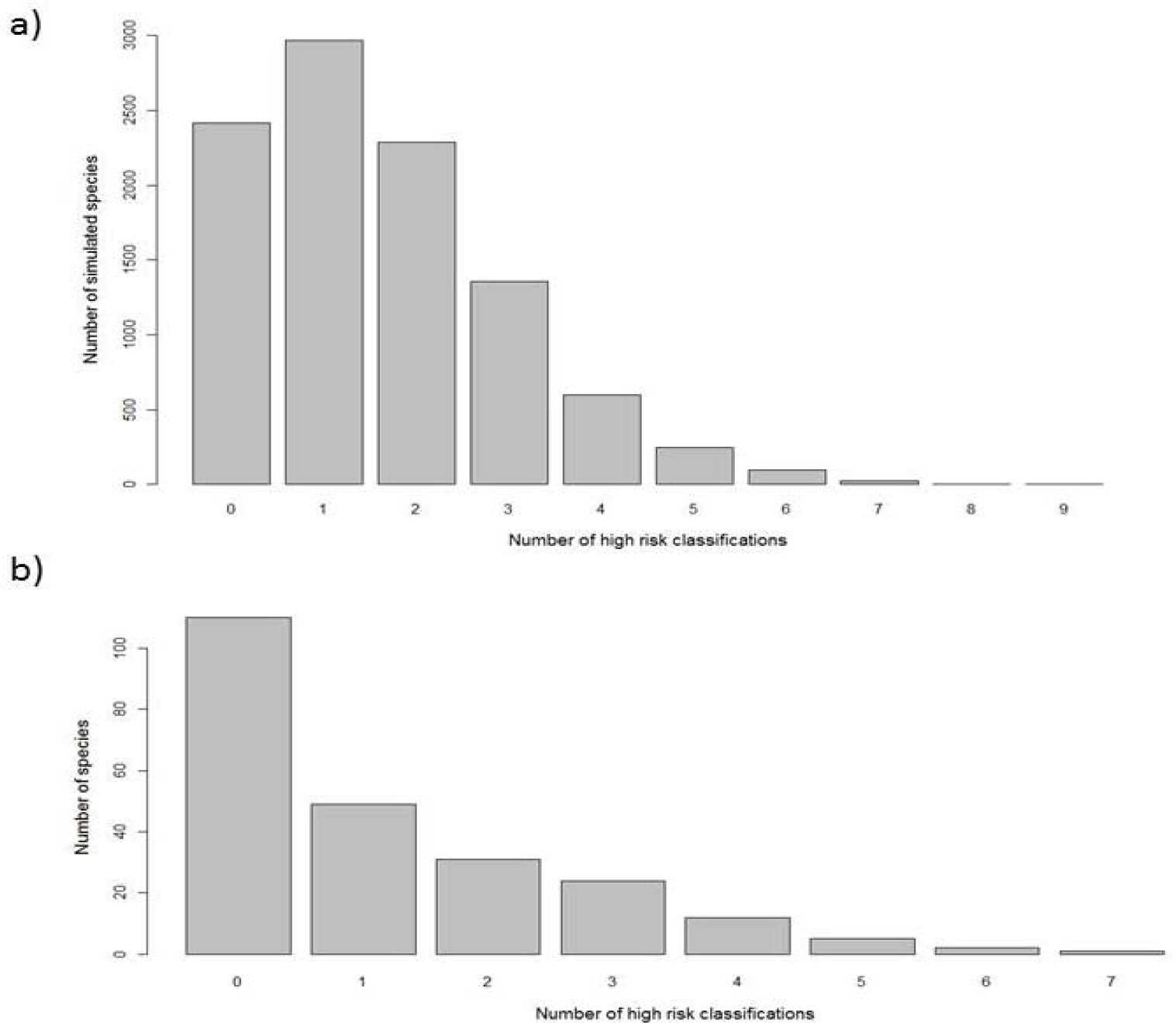
Frequency distribution of high risk classifications for a} simulated species and b).real species assessed with historic data. The number of risk assessment frameworks under which each simulated or real species was classified as high risk.

Sufficient data to run all the frameworks are only available for a small subset of taxonomic groups (primarily vertebrates, and birds in particular), which only samples a relatively small range of potential species-types, and hence of ecological traits. In order to sample the full range of potential trait variation in nature, we generated 10,000 ‘simulated species’, each with randomly generated trait sets and populations, bounded by real world (trait value) limits. To fully incorporate all possible parameter space, we chose to remove all but logically necessary correlations between traits (e.g. we retained logical consistency between numbers of habitats occupied and presence in particular habitat types, but did not enforce correlations between body size and fecundity, which are positive in some taxa but negative in others). Correlations between life history traits vary widely between taxonomic groups and would be almost impossible to simulate accurately for a wide range of taxa. As our extensive real bird dataset maintains correlations between traits for that group, our simulation provides contrasting data by removing such constraints, increasing the generality of our assessment.

All 10,000 simulated species were assessed individually using each of the 12 risk assessments. The frameworks show broadly similar patterns in the overall assignment of risk to the real species, classifying the majority of species as low risk and relatively few as high risk (Supplementary Figure 1). However, over 75% of the 10,000 simulated species were classified as high risk by at least one framework considered, and only 135 were assessed as high risk by more than half of the frameworks (Figure 1a). Overall, we found poor agreement across the frameworks in assigning risk (Figure 2, r_s_ mean = 0.07 ± 0.01, r_s_ median = 0.04). Pairwise correlations within broad framework types were stronger than the overall pairwise correlations (between trait-based frameworks: r_s_ mean = 0.13 ± 0.04, r_s_ median = 0.08; between trend-based frameworks: r_s_ mean = 0.29 ± 0.12, r_s_ median = 0.18), but still relatively poor. There was also little difference between frameworks designed for single species and more generic frameworks (between species-specific frameworks: r_s_ mean = 0.09 ± 0.05, r_s_ median = 0.04 and between generic frameworks: r_s_ mean = 0.11 ± 0.03, r_s_ median = 0.04). Using inter-rater reliability analysis to compare agreement between frameworks in their classification of simulated species in the highest risk category only, we again found weak overall agreement (mean κ_PABAK_ = 0.55 ± 0.02, median κ_PABAK_ = 0.52). This inconsistency suggests against using a consensus of contrasting methods as the basis for prioritisation.

**Figure 2.**
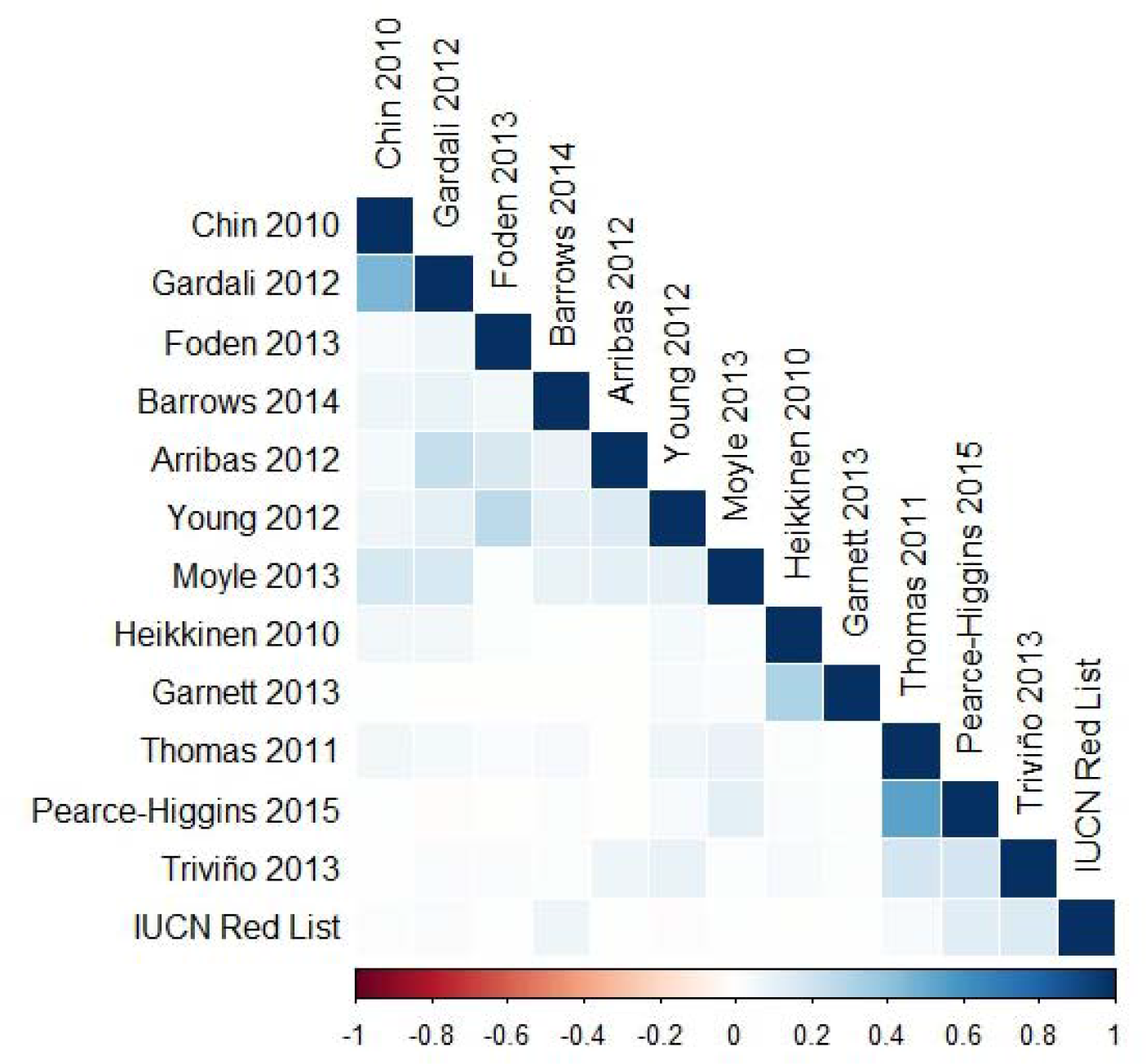
Correlation matrix showing spearman rank correlation coefficients (r_s_) for each of the 12 frameworks, pairwise against the others and the Red List outputs for the simulated species. The matrix is a visual representation of the r_s_ value (see x axis for range), with darker blue indicating a stronger positive correlation; using output data for the 10,000 simulated species. The correlations between each of the climate change risk assessment frameworks and the simulated Red List risk category are shown in the bottom row of the matrix. Reference numbers are as in Table 1.

Comparing the outputs of the frameworks to Red List outputs also produced poor correlations (Figure 2: Spearman’s rank correlation r_s_ mean = 0.04 ± 0.01, r_s_ median = 0.01), with trait-based assessments showing weaker correlation with Red List outputs than trend-based approach types (trait based: r_s_ mean = 0.02 ± 0.01, r_s_ median = 0.01, trend based: r_s_ mean = 0.11 ± 0.01, r_s_ median = 0.13).

To investigate similarities between the risk assignments of different frameworks further, we used Principal Components Analysis (PCA) on the risk category outputs. We found distinct clusters for trait-only frameworks^10–14^ and trend-based frameworks^15–17^ with hybrid assessments falling between the two^18, 19^ (Figure 3, Supplementary Table 5). This pattern is the same for the pairwise correlations between frameworks, with weak agreement overall, but stronger correlations within the five purely trait-based frameworks and within the three trend-based frameworks.

**Figure 3.**
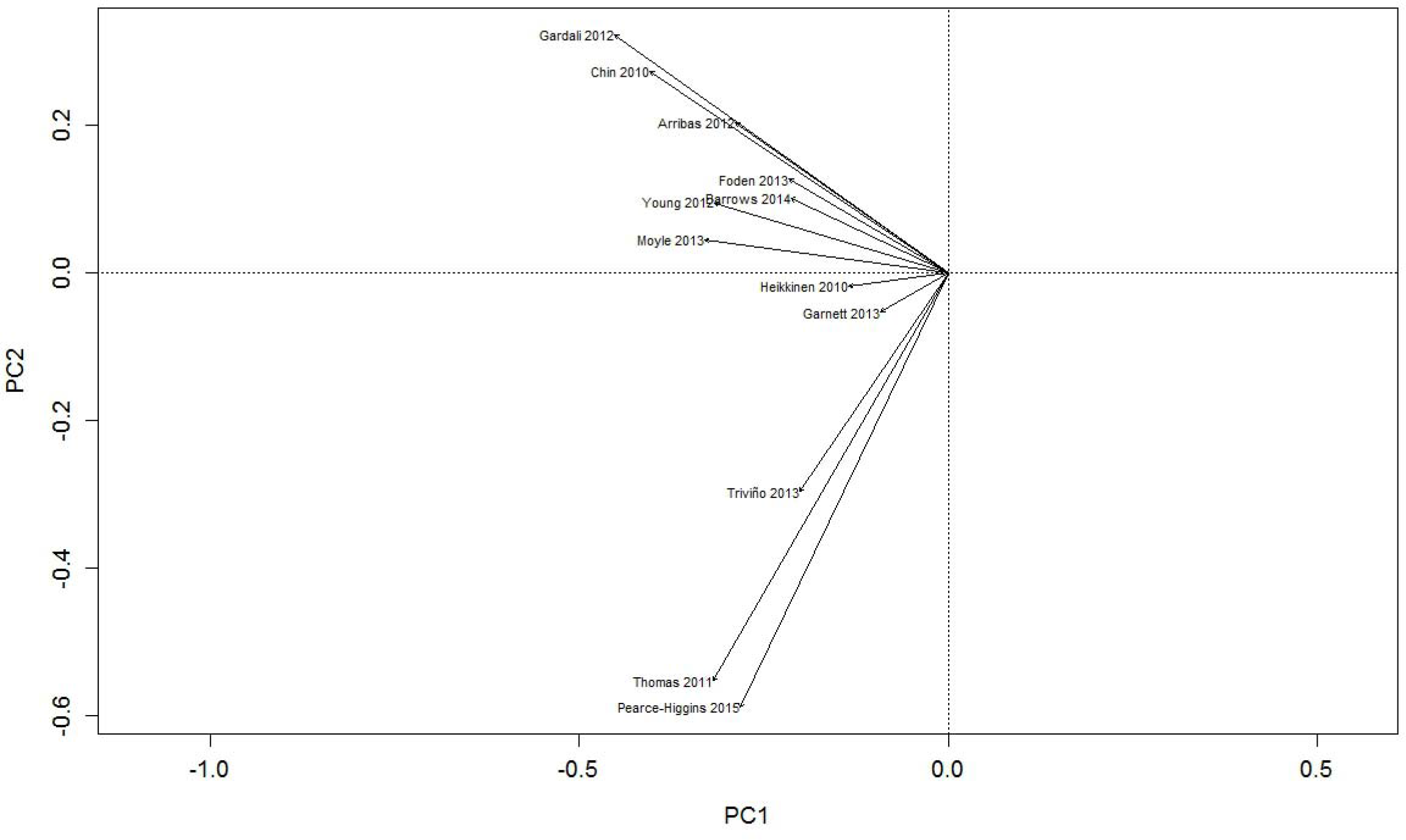
Principal component biplot. The first two principal components obtained by applying principal components analysis to the risk category outputs from the 12 frameworks for the 10,000 simulated species. Reference numbers are as in Table 1.

### Validation of different vulnerability frameworks

Given the great variation in the risk categories assigned to each real and simulated species, validation is required to assess whether any of the vulnerability frameworks has any predictive power at all. To do this, we used historic data for British birds and butterflies to assign each species to a risk category (for each of the 12 frameworks), using data up to the 1990s. We then used 1990s-2000s observed trends in distribution/abundance to evaluate whether the risk categories assigned by each framework were predictors of subsequent population and distribution changes. We ran the assessments for 169 British bird species (validated against observed distribution and abundance changes) and 50 British butterfly species (validated against abundance changes only), for which data were available to model distribution change under a medium emissions climate change scenario and to calculate recent changes in distribution/population. The risk outputs of each framework were again standardised using the same Low/Medium/High scale defined previously (Supplementary Table 2). Because species are affected by multiple factors in addition to climate change (e.g. a low-risk species may decline for non-climatic reasons), we used quantile regression to consider trends in distribution/population change in the 0.50 and 0.75 quantiles, representing the response of the majority of species within each risk category.

Overall, none of the frameworks showed strong predictive power, with 8 of the 12 showing no significant trend in either the 0.50 or 0.75 quantiles when comparing risk category against observed change in distribution for the British birds (Figure 4). Of the remaining four assessment frameworks, two^13, 20^ showed significantly worse-than-random risk categorisations – higher risk species showed more positive subsequent distribution trends than lower risk species (the two frameworks that show a significant positive trend for the 0.75 quantile, in Figure 4). Only two^15, 17^ of the frameworks produced significantly better-than-random risk assessments (one significant for the 0.50 and 0.75 quantiles, and one for the 0.75 quantile). Both of these frameworks are trend-based approaches, which would suggest incorporating this type of data into the assessment process produces more robust risk outputs.

**Figure 4.**
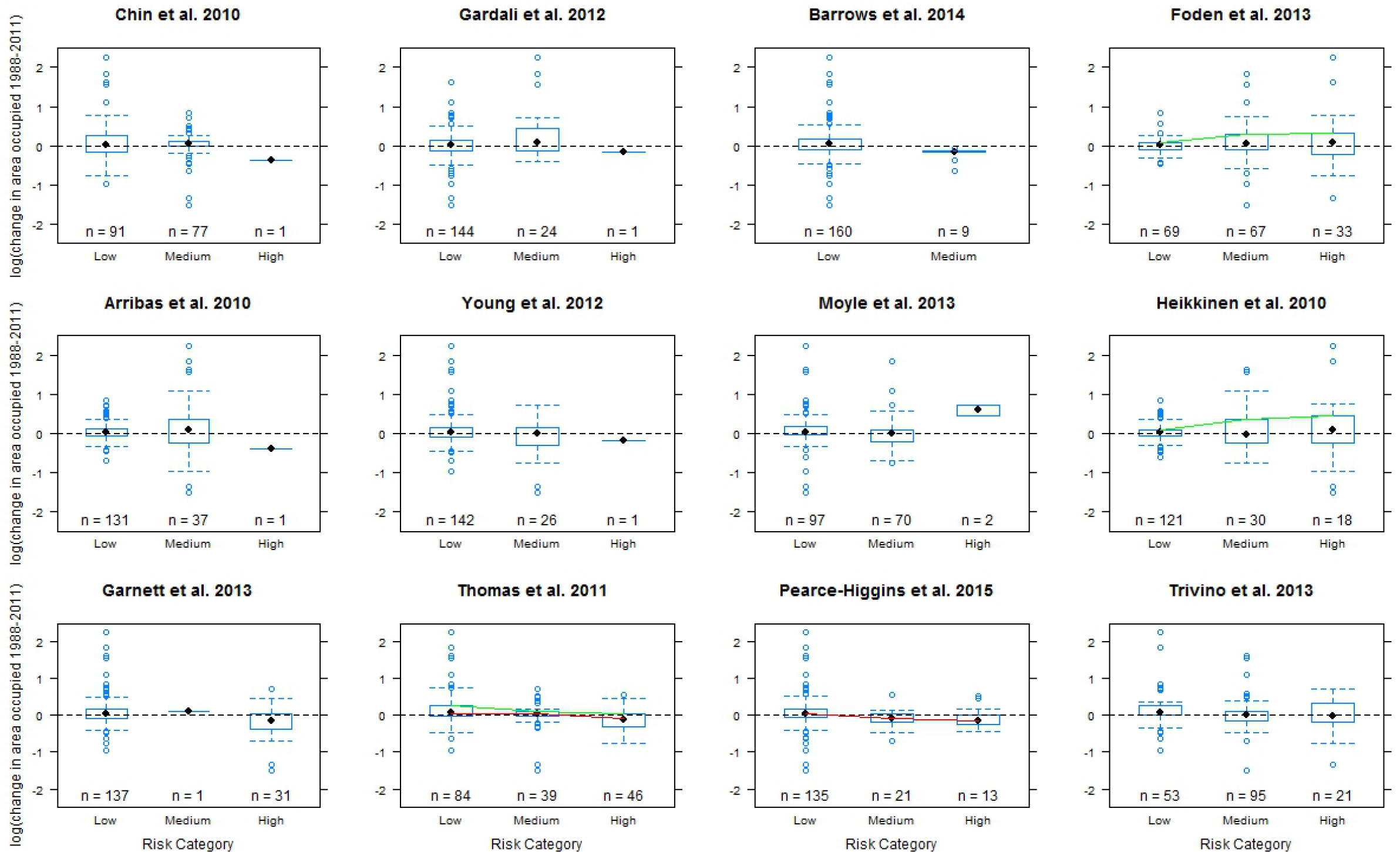
Validation boxplots showing logged change in bird distribution against simplified risk category for each of the 12 risk assessment frameworks. Red lines show a significant trend in the 0.50 quantile and green lines show a significant trend in the 0.75 quantile. Assessments are for 218 British bird and butterfly species.

The results of validation for both birds and butterflies when using population change, rather than distribution change as the response variable, also suggested limited framework effectiveness. When considering changes in bird populations, there were no significant trends in the 0.50 quantile for any of the frameworks and only a single framework showed a significant trend for the 0.75 quantile (Supplementary Figure 2). This framework^20^ provided significantly worse-than-random risk categorisations (high risk species subsequently showed greater population growth), and was one of the two frameworks that was also worse than random when assessed against bird distribution changes. There were no significant trends in either the 0.50 or 0.75 quantile for any of the 12 frameworks when assessing population change for butterflies (Supplementary Figure 3), although overall performance appeared to be better than for the bird population analysis.

Frameworks are ranked in Table 3 first by significant ‘correct’ predictions (high risk species subsequently decline most) across the six tests (0.50 and 0.75 quantiles for each of bird distributions, bird abundances, butterfly abundances), and then by ‘correct’ non-significant trends. Our validation tests therefore suggest a few of the frameworks may work but that others have no predictive power and some are worse than random, given the test data.

### Validation using an ensemble approach

In addition to the individual framework validation, we also consider the effectiveness of using an ensemble approach to climate vulnerability assessment. We compared the modal risk category assigned to a species by the 12 frameworks against the same change in distribution/population value used in the individual framework validations. For the 169 bird species, only two had a modal risk classification of high risk, with both showing positive changes in distribution (Figure 5a) and population (Figure 5b), measured over the validation period. The 50 butterfly species also had just two species with a modal high risk classification, with one increasing its population over the validation period and the other showing little change in its population (Figure 5c). Therefore, the ensemble approach did not identify high risk species that subsequently declined – and across all species, there was no link between the consensus risk category and subsequent distribution trends in quantile regressions. We also considered the maximum risk category assigned by an ensemble approach (Supplementary Figure 4), but as this approach was not significant either; and it would be impractical to use to set conservation priorities because maximum risk identified over half the bird and butterfly species as high risk (Figure 1b).

**Figure 5.**
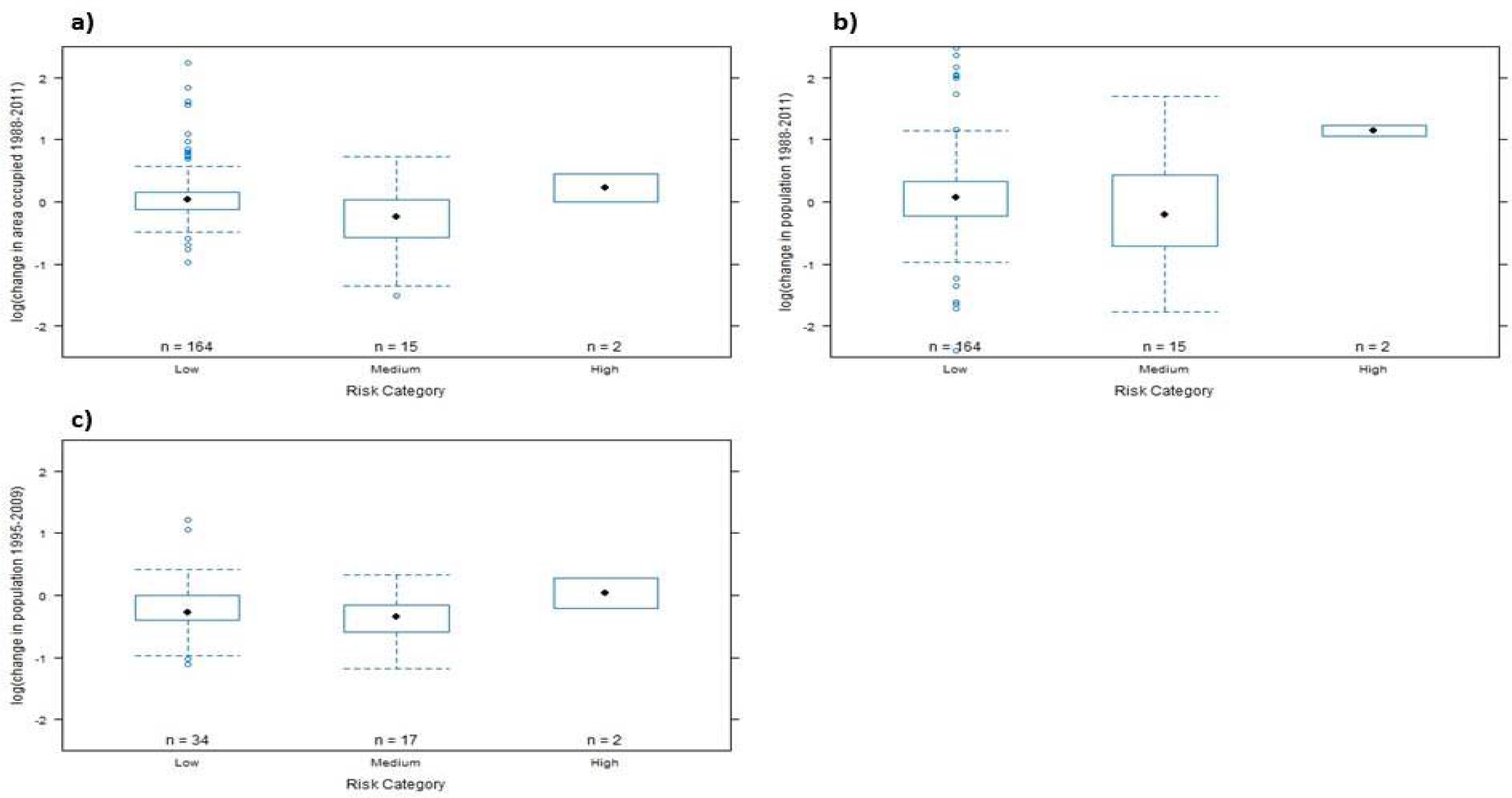
Validation boxplots showing a) logged change in bird distribution, b) logged change in bird population and c) logged change in butterfly population, against modal simplified risk category from across all 12 risk assessment frameworks.

## Discussion

Our results from both real and simulated species highlight poor overall agreement on risk assessment across the 12 frameworks considered, particularly between trend-and trait-based approach types, suggesting that the differences between approach types are fundamental. More importantly, our validation results suggest that few methods have any predictive value, at least for the test-data considered. The inconsistencies between methods holds, regardless of whether we take into account the correlated traits that exist for real species within a given taxonomic group or if we minimise correlations between traits in simulated species (given that different higher taxa possess dissimilar trait correlations). The similarities between our results for simulated and real species suggests that the inconsistencies arise from differences between the risk framework methods themselves (i.e., which variables are included in an assessment, and how they are combined to place each species in a risk category) rather than from the test data that we used. Given that real and simulated species are placed in different climate-risk categories by different risk assessment frameworks, it is essential that validations are carried out to assess whether none, some or all of the frameworks have predictive power.

The results from the validation analysis revealed that most frameworks perform poorly. Across the six validation tests (0.50 and 0.75 quantiles for each of bird distribution, bird abundance and butterfly abundance changes), two frameworks^13, 20^ produced significantly worse-than expected assessments in one or more cases, in the sense that the species assigned to high-risk categories subsequently showed more favourable distribution and/or population trends than the species that were assigned to lower risk categories. Another three frameworks^10,12,19^ gave qualitatively (though not significantly) similar results; i.e., the bottom five frameworks in Table 3 performed poorly. This leaves the ‘top seven’ for further consideration. Of these, only two methods^15, 17^, both of which were trend-based, assigned risk appropriately (i.e. the high-risk species declined more than lower risk species) and significantly (Figure 4); although predictions were only significant when considering change in distribution as the response variable, not change in population (top two rows of Table 3). One of these methods^15^ also generated non-significant predictions in the expected direction in all of the other tests (top row of Table 3). These two methods are closely related to one another, with both using predicted trends based on climate as the driving force, with one^15^ using additional trait/habitat information that modifies the capacity of each species to respond as predicted. These additional constraints apparently increased the predictive power of this framework.

**Table 3.** Summary validation trends. The direction of the trend in either distribution or abundance change for birds and butterflies from Low risk species to high risk species, with a negative trend indicating the framework is performing as expected and a positive trend indicating poor framework performance. Significant trends are denoted with *. The frameworks are ranked first by number of significant negative trends and then by number of non-significant negative trends.

| Framework | Methodology Type | Bird distribution trend direction |  | Bird population trend direction |  | Butterfly population trend direction |  | Correct significant trends | Correct non-significant trends | Rank |
| --- | --- | --- | --- | --- | --- | --- | --- | --- | --- | --- |
|  |  | 0.50 quantile | 0.75 quantile | 0.50 quantile | 0.75 quantile | 0.50 quantile | 0.75 quantile |  |  |  |
| Thomas et al. 2011<sup>15</sup> | Trend | -* | -* | - | - | - | - | 2 | 4 | 1 |
| Pearce-Higgins et al. 2015<sup>17</sup> | Trend | - | -* | - | + | - | + | 1 | 3 | 2 |
| Young et al. 2012<sup>18</sup> | Hybrid | - | - | - | - | - | - | 0 | 6 | 3.5 |
| Barrows et al. 2014<sup>14</sup> | Trait | - | - | - | - | - | - | 0 | 6 | 3.5 |
| Garnett et al. 2013<sup>21</sup> | Hybrid | - | - | - | + | - | - | 0 | 5 | 5 |
| Arribas et al. 2012<sup>11</sup> | Trait | - | - | + | + | - | - | 0 | 4 | 6.5 |
| Triviño et al. 2013<sup>16</sup> | Trend | - | + | - | - | - | + | 0 | 4 | 6.5 |
| Gardali et al. 2012<sup>12</sup> | Trait | - | - | + | - | + | + | 0 | 3 | 8.5 |
| Chin et al. 2010<sup>10</sup> | Trait | - | - | + | - | + | + | 0 | 3 | 8.5 |
| Moyle et al. 2013<sup>19</sup> | Hybrid | + | + | + | + | - | - | 0 | 2 | 10 |
| Foden et al. 2013<sup>13</sup> | Trait | + | +* | + | + | - | - | 0 | 2 | 11 |
| Heikkinen et al. 2010<sup>20</sup> | Hybrid | + | +* | + | +* | + | + | 0 | 0 | 12 |

Some of the other frameworks do show a similar overall pattern, but assign such small numbers of species to the high risk category that it was not possible to detect significant trends (see Figure 4). For example, one trait-based framework^14, 18^ failed to assign any species to the high-risk category (and only between 9 and 13 to the medium-risk category) and one hybrid framework^18^ only assigned 1, 1 and 5 species to high risk in the three tests. Two of the frameworks^20, 21^ classify species into risk categories based on proportions (e.g. top tenth of values assigned high risk) instead of consistently set threshold values, as seen in the other frameworks. The risk outputs from these two frameworks correlate poorly with most others, and they fall close to the origin in the PCA (Figure 3). Another framework^13^ uses proportional cut offs for some input data and along with a method that uses proportional risk categories^20^ performs poorly overall in the validation analysis; with significant trends in the opposite direction to that expected if assigning risk suitably. Proportions of species at risk from climate change are not expected to be the same in different regions (or taxonomic groups), so we recommend avoiding proportional approaches.

Since each framework we tested gives markedly different results, it is necessarily the case that most or all methods are misleading, which limits informed conservation responses. A possible alternative is to consider the results from an ensemble of climate vulnerability assessments. The high variability in outputs, however, also limits the effectiveness of taking an ensemble approach. We considered two possible approaches to this. The first was to consider the possibility that there are many different mechanisms of endangerment from climate change, and hence to consider a species as at risk if any of the 12 methods classified it as at high risk. This was not practically useful because the majority of species were identified as high risk using this approach. The second was to assign species to the modal class of vulnerability, which resulted in almost no species being classified as high risk. Neither approach significantly identified declining species in validation. The output from the ensemble of methods does not offer sufficient improvement over any individual method to justify the time and effort required to collect the data to run all of the assessments.

It should be noted that the time period for the observed changes used in the validation analysis are relatively short for both birds and butterflies (20 and 10 years respectively), and from a period when a range of other pressures have also affected species’ population in the area considered, particularly changes in agricultural management^26^. There is a possibility that some species considered may be climate-threatened but not yet showing a strong negative response in distribution or population, whilst others may be limited by other factors, potentially leading to the under-estimation of framework performance. However, we would expect frameworks to show some separation between range- or population-expanding and contracting species, as during this period both bird and butterfly communities have responded to climate change^27, 28^, for example with polewards shifts^29–31^. The fact we do not see such a pattern for most assessments (and some trends are the reverse of those expected), combined with the results of our comparison between frameworks, does highlight the lack of evidence currently available to support the use of most of these frameworks. As some of the assessments are designed for global assessments of risk, there is a possibility that the poor performance is a consequence of applying them over a regional scale. However, this methodology is being applied at non-global scales by researchers and practitioners^32^ so the results of our validation at a regional scale remain applicable to how the methods are actually being used.

The science underpinning trend-based approaches is stronger; with increasing evidence that species distribution models used to measure exposure in trend-based approaches can retrodict recent population and range trends^33–35^. There remains uncertainty around identifying the key traits influencing species vulnerability to climate change^24^, which may vary widely by taxonomic group and could explain the wide range of inputs across the different trait-based assessments. Recent work^36^ has advocated the combination of elements of trait-based vulnerability assessments with species distribution modelling to produce more realistic projections of future risk. This approach has already been implemented to different extents by some frameworks considered here^15,16,18^, although the outputs of these show at best weak correlations with purely trait-based assessments, suggesting that trait-only assessments may not adequately capture the exposure component of climate risk. The two general types of assessment (trait, trend) effectively represent different paradigms, with combined approaches representing arbitrarily-weighted blends of the two.

We have demonstrated that different vulnerability assessment frameworks should not be used interchangeably when attempting to assess a species’ potential future risk to climate change, because assessments made with either real or simulated species produce conflicting results. Our validation results suggest there is currently little evidence to support the use of purely trait-based vulnerability assessments. Trend-based approaches are the only type of methodology to consistently and significantly assign species to appropriate risk categories in the validation analysis, particularly when this information is supplemented with additional species trait data. Whilst we recognise this may restrict the assessment options available to practitioners (e.g. without long-term monitoring data, trend-based approaches will not be possible), our results highlight the considerable uncertainty in the results of approaches not incorporating this type of information. A poorly performing framework should not be used simply because it is the only one for which adequate data are available. Without significant investment in long-term monitoring, to study change as it occurs, and in research to identify exactly what traits make a species’ vulnerable to climate change, our ability to identify the species most in need of conservation attention in the face of climate change will remain poor.

## Methods

### Exemplar and real species comparisons

The assessments of exemplar real species and additional British bird species (Table 2) were carried out based on trait and distribution data within Great Britain, due to the quality and availability of data for the taxa considered within this region. The 18 exemplar species were chosen because they were the only species of any taxonomic group with both comprehensive distribution (in two or more time periods) and traits data and a northern or southern range margin lying within Great Britain^37^ (species with range boundaries in a region are likely to be of interest when running climate change vulnerability assessments). All common British breeding bird and butterfly species were considered for the additional assessment, the 218 species selected being the ones for which future distributions could be modelled based on data availability.

Trait data for the real species were collected from a variety of sources including scientific literature and species atlas data^38, 39^. Projected distribution changes were based on existing bioclimate model data^17^, applying a Bayesian, spatially explicit (Conditional Autoregressive) GAM^40^ to the bird and butterfly distribution data. A medium emissions scenario (UKCP09 A1B) for projected climate change for 2080 was used for future climate data, corresponding to a 4°C increase in average temperature. The assessments were also run using a low emissions scenario (UKCP09 B1), corresponding to a 2°C increase in average temperature, with little difference in overall risk category assignment (Supplementary Table 4).

### Simulated species comparisons

To compare the outputs of the 12 risk assessment frameworks using simulated species, we generated ranges of values for the 117 unique input variables (Supplementary Table 1), covering characteristics such as species traits and population trends. We then drew values for each of these input variables to generate 10,000 combinations of ‘trait sets’ that were used as simulated species in the assessments, in lieu of real world data for many species. Where it has been possible to do so, we applied constraints on the input variables to ensure logical consistency. For example, in the case of interspecific interactions, some frameworks ask broadly whether there is a dependence of a species on any interspecific interaction, whilst other frameworks require inputs relating to multiple, clearly-defined interspecific interactions. In this situation it would not make sense for the broad interaction to be scored as absent while specific interactions are scored as present. In this case the broad interaction is generated first and the scores of more specific interaction variables are influenced by that, to ensure consistent inputs across frameworks.

For continuously distributed input variables, upper and lower bounds were set based on reported values from the literature (e.g. body size, generation time) or theoretical minimum and maximum values. A value for the variable for each simulated species was then drawn from a uniform distribution bounded by those upper and lower limits. Species current distributions were simulated using the same approach, sampling a value for area occupied (in km^2^) from a uniform distribution with an upper limited based on known real world distribution limits. For projected changes to species distributions under climate change, a future distribution was generated using the same process as for current distributions, and the percentage change in area between the two calculated.

The uniform distribution was chosen for all variables (equal probability for binary and categorical variables) because, for many input variables, there was little or no data available on how they might be distributed in reality (and they differ greatly between taxonomic groups), so an arbitrary selection of distribution would have been needed. Nonetheless, where there was an a priori expectation of the distribution of a trait based on the literature (e.g. logarithmic scaling of dispersal distance), the uniform draw was from between the transformed trait limits. The uniform distribution also allows for generation of traits covering the full range of the potential parameter space for the input variables, which was one of the main advantages of generated trait sets rather than a larger sample of real species data. The results therefore test consistency in framework performances, rather than the ‘true’ frequencies of risk (which we do not know, given the differences between framework methods).

Many of the input variables are categorical, typically scored as low/medium/high or some similar variation. In some cases it is possible to base these on a continuous variable which is then split into the different categories (e.g. dispersal distance < 1km scored as low, dispersal distance > 1km and < 10km scored as medium, dispersal distance > 10km scored as high). Where it has not been possible to generate a continuous variable to base the categorical split on (e.g. impact of climate mitigation measures – scored as low to high), the category was instead assigned randomly to one of the possible options, with an equal probability of assignment to each. IUCN Red List conservation status was required as an input to one of the frameworks and was generated using IUCN criteria A to D, with no projected future changes considered. This conservation status for each simulated species was also used in comparisons of Red List risk category against risk category for each framework, and therefore informs us of the relationship between climatic and non-climatic risks rather than whether the Red List could adequately take climate change into account.

### Validation

To examine how well the different climate vulnerability assessments performed at projecting future risk we used the results of assessments based on historic species data to compare against observed recent trends in species distribution/abundance. For validation of the frameworks to produce robust results they need to be tested using reliable input data, poor quality input data will always lead to poor assessments of risk regardless of the method used for the assessment. We therefore utilized some of the best quality data available globally and selected British birds and butterflies for the analysis.

Validations were carried out by using historically-available data to assign species to low-, medium-and high-risk categories (for each of the 12 risk assessment frameworks), as though the assessments were carried out in the past, and then we compared recent distribution and population changes for species that had been assigned to each risk category. Assessments for British birds were based on the time period 1988-1991, to match the breeding bird atlas data^41^. Assessment inputs based on the ‘then-current’ distribution/population were calculated from this Atlas data, with historic changes in distribution calculated from the 1968-1972 Atlas to the 1988-1991 Atlas^41^. Projected changes in distribution were modelled using the 1988-1991 Atlas distribution data and future climate projections for 2080. Historic assessments for British butterflies were performed using the same approach, based on the 1995-99 Millennium Butterfly Atlas^39^ and historic trends calculated from the previous 1970-82 national survey. Future projected distributions were modelled using the same methodology as for the bird species.

In addition to the output of the assessments, observed recent trend data for distribution and population change since the assessment time period was required. For bird distribution trends, data from the 2007-2011 Atlas was used giving the percentage change in occupied 10km grid squares between 1988-1991 and 2007-2011. Observed changes in population for birds were obtained from the State of the UK Birds report^42^ as a percentage change in population from 1995 to 2013. Butterfly population change data was obtained from the State of the UK Butterflies report^43^, giving a percentage change in population from 1995 to 2005. Although these dates partly overlap with the Millennium Butterfly Atlas^39^, the population data are collected on fixed transects that are separate from the millions of independent distribution records that give rise to the Atlas maps. Distribution change data for the butterflies was not used in the analysis due to a large increase in observer effort in latter time period, which resulted in increases in distribution that are likely to reflect increased effort rather than true changes in distribution.

### Statistical analysis

The risk category outputs from each of the frameworks were converted to a set of standardised categories: Low/Medium/High risk (Supplementary Table 2). Broad agreement between the frameworks was tested on a pairwise basis using Spearman’s rank correlation, to establish how consistently species were assigned to the same Low/Medium/High risk categories by the different frameworks.

Rank correlation allows for a comparison of how well the different frameworks correspond across all levels of risk assignment, but a potentially more useful comparison is of how well they agree in identifying a species as high risk, based on the assumption that assessments will primarily be run to identify the species most vulnerable to climate change. To compare agreement on just high risk species, the risk categories were further simplified to a binary, ‘low and medium’ versus ‘high’ categorisation. Cohen’s kappa, a measure of inter-rater reliability, was calculated to compare agreement between frameworks. The prevalence and bias-adjusted Cohn’s kappa (PABAK)^44^ was used due to the relatively low frequency of species scoring as high risk.

Principal component analysis (PCA) was used to examine how much of the variation in risk assignment was influenced by certain frameworks and to identify whether frameworks of the same general type (trait, trend) showed similar patterns in risk category assignment. Risk category outputs from each framework for the 10,000 simulated species were used in this analysis.

We predicted that all species at high risk due to climate change should have seen population/distribution decreases, whilst species identified as low risk may have increased, decreased or not changed their population/distribution if factors other than climate are driving the changes. We therefore used quantile regression to validate framework performance, with change in distribution or abundance as the response variable and framework risk categorisation (Low/Medium/High) as the predictive factor^45^. This allowed us to consider trends in the upper quartiles of distribution/population change instead of just the mean, which would identify if the majority of high risk species are declining as we would expect if a framework is performing well. Both the 0.50 and 0.75 quantiles were considered in the analysis, and the models were tested for significance against a null model using an ANOVA.

## Author Contributions

All authors conceived and designed the study; CJW, CMB, CDT and RC collected data; CJW and CMB analysed the data; all authors interpreted the results; CJW produced the original draft and all authors contributed to revisions

